# Mutation Patterns Predict Drug Sensitivity in Acute Myeloid Leukemia

**DOI:** 10.1101/2023.05.24.541944

**Authors:** Guangrong Qin, Jin Dai, Sylvia Chien, Timothy J. Martins, Brenda Loera, Quy Nguyen, Melanie L. Oakes, Bahar Tercan, Boris Aguilar, Lauren Hagen, Jeannine McCune, Richard Gelinas, Raymond J. Monnat, Ilya Shmulevich, Pamela S. Becker

## Abstract

Acute myeloid leukemia (AML) is an aggressive malignancy of myeloid progenitor cells characterized by successive acquisition of genetic alterations. This inherent heterogeneity poses challenges in the development of precise and effective therapies. To gain insights into the genetic influence on drug response and optimize treatment selection, we performed targeted sequencing, *ex vivo* drug screening, and single-cell genomic profiling on leukemia cell samples derived from AML patients. We detected genetic signatures associated with sensitivity or resistance to specific agents. By integrating large public datasets, we discovered statistical patterns of co-occurring and mutually exclusive mutations in AML. The application of single-cell genomic sequencing unveiled the co-occurrence of variants at the individual cell level, highlighting the presence of distinct sub- clones within AML patients. Machine learning models were built to predict *ex vivo* drug sensitivity using the genetic variants. Notably, these models demonstrated high accuracy in predicting sensitivity to some drugs, such as MEK inhibitors. Our study provides valuable resources for characterizing AML patients and predicting drug sensitivity, emphasizing the significance of considering subclonal distribution in drug response prediction. These findings provide a foundation for advancing precision medicine in AML. By tailoring treatment based on individual genetic profiles and functional testing, as well as accounting for the presence of subclones, we envision a future of improved therapeutic strategies for AML patients.

**One Sentence Summary:** Integrative computational and experimental analysis of mutation patterns and drug responses provide biologic insight and therapeutic guidance for patients with adult AML.

## Introduction

Acute myeloid leukemia (AML) is a hematologic malignancy characterized by maturation arrest and uncontrolled proliferation of myeloid progenitor cells (Siveen et al., 2017). Data from several large cohorts of AML patients have been analyzed to understand the mutational landscape and how the genetic diversity of the cancer defines the pathophysiology of AML (Cancer Genome Atlas Research et al., 2013; Gerstung et al., 2017; Papaemmanuil et al., 2016; Tyner et al., 2018). Characterization of the gene mutations from different studies yields a highly complex pattern of potential driver events for AML. According to bulk molecular profiling, the acquisition of mutations in leukaemogenesis follows a stepwise pattern, where mutants with high variant allele frequencies emerge early, while mutations with lower variant allele frequencies are thought to occur later (Genovese et al., 2014; Jan et al., 2012). Single-cell mutational profiling showed AML is dominated by a small number of clones, which frequently harbor co-occurring mutations in epigenetic regulators (Miles et al., 2020). Mutations in signaling genes often occur in distinct subclones from the same patient (Miles et al., 2020). The complexity of malignant cell evolution and presence of heterogeneous sub-clones make it challenging to stratify patients and optimize treatment.

There are only a limited number of FDA approved targeted inhibitors for AML, such as the FLT3 inhibitors midostaurin and gilteritinib, the IDH1 inhibitor ivosidenib, the IDH2 inhibitor enasidenib, tyrosine kinase inhibitors for KIT mutations, and the BCL2 inhibitor venetoclax. There are several newer agents under investigation, but the overall number of such inhibitors remains relatively small compared to the known number of mutated genes. Novel methods for inhibiting mutated genes, such as targeted protein degradation (Khan et al., 2019) and antibodies to drug-peptide complexes (Zhang et al., 2022), appear promising, but remain under development. Thus, the path to optimize individual treatment remains complex with inadequate options.

Different factors may affect drug sensitivity, such as the genetic alterations, subclonal evolution, the phenotypic states of cells and the bone marrow microenvironment that includes the immune response. Nevertheless, the majority of patients lack the genetic mutations targeted by the approved drugs. Co-occurrence of gene mutations has been widely reported in many cancer types, including AML (Papaemmanuil et al., 2016). It has been reported that leukemia stemness and co-occurring mutations drive resistance to IDH inhibitors in AML (Wang et al., 2021). The co-occurrence of gene mutations can be due to clonal evolution with acquisition of additional mutations in the same cell or to distinct new clones with other mutations (Miles et al., 2020), which may require different treatments or their combinations to optimize response.

Several large projects have characterized the genomic features of AML, such as the TCGA-LAML project that characterized genomic alterations for 200 patients using Whole Exome Sequencing (WES) or Whole Genome Sequencing (WGS) (Cancer Genome Atlas Research et al., 2013), the Beat AML project that characterized the genomic features for 632 AML patients using WES (Tyner et al., 2018), and the German-Austrian AML Study Group (AMLSG) that utilized targeted sequencing in 1540 AML patients (Gerstung et al., 2017; Papaemmanuil et al., 2016). Additionally, *ex vivo* drug screening has been carried out in the Beat AML dataset, with a set of 125 drugs(Tyner et al., 2018) as well as in the FPMTB study in Finland that performed *ex vivo* drug sensitivity assays for 515 anti-cancer drugs (Malani et al., 2022). These publicly available datasets allow us to explore the genetic heterogeneity of AML cells and predict drug sensitivity for patients with given genetic alterations. However, more drugs are needed to treat AML patients with clear genetic alteration patterns for which there are still no specific inhibitors.

In this study, we sought to characterize the heterogeneous genotypes of AML patients with different co-occurring or mutually exclusive mutation patterns and to associate such patterns with drug response to develop clinical prediction models. Using the abundant public data resources for AML, we have built co-mutation graphs and identified sub-graphs that allow us to understand the potential functional impact of different mutation patterns and predict drug sensitivity. To validate the relationships identified by the analysis of public data, we also characterized the mutational landscape of AML patients using targeted sequencing of 194 AML-related genes and carried out *ex vivo* drug screening using a library of 208 drugs for 99 patients. Using the genetic analysis of these patients, we identified potentially effective drugs for patients with various genetic backgrounds. We also performed single cell genomic profiling to confirm the co-occurrence of mutations at the single cell level and constructed machine learning models for the prediction of drug sensitivity using the genetic variants. Our newly generated targeted sequencing, *ex vivo* drug screening, and single cell genomic profiling datasets, together with the accompanying analysis, provide options for potentially effective selection of therapies.

## Results

### Dataset overview

This study includes a cohort of 99 patients with AML of which >70% had relapsed, with a median age of 58 (range 19-83). The distributions of clinical features such as antecedent hematologic disorder (AHD), European LeukemiaNet (ELN2017) classification, complete remission (CR), duration of overall survival, and the status of patients (de novo or relapse) are shown in Figure 1A (data available in Supplementary Table 1 and 2). Myeloid blasts were enriched to >80% prior to NGS or high throughput drug screening using magnetic beads when needed. The enriched blasts from the AML patients were subjected to *ex vivo* drug screening and targeted sequencing. We collected drug screening data for all 99 samples and full targeted sequencing data for 75 out of the 99 samples (Figure 1B) owing to the limitation in cell numbers. With the targeted sequencing and *ex vivo* drug screening datasets, we then analyzed the mutation patterns for AML patients using inhouse mutation data and public datasets, built machine learning models to predict drug sensitivity for AML patient derived samples, and identified the features that are predictive of sensitivity and resistance (Figure 1B). Single cell genomic sequencing for 8 samples was performed to confirm the mutation pattern at the single cell level and reveal mutational heterogeneity.

**Figure 1.**
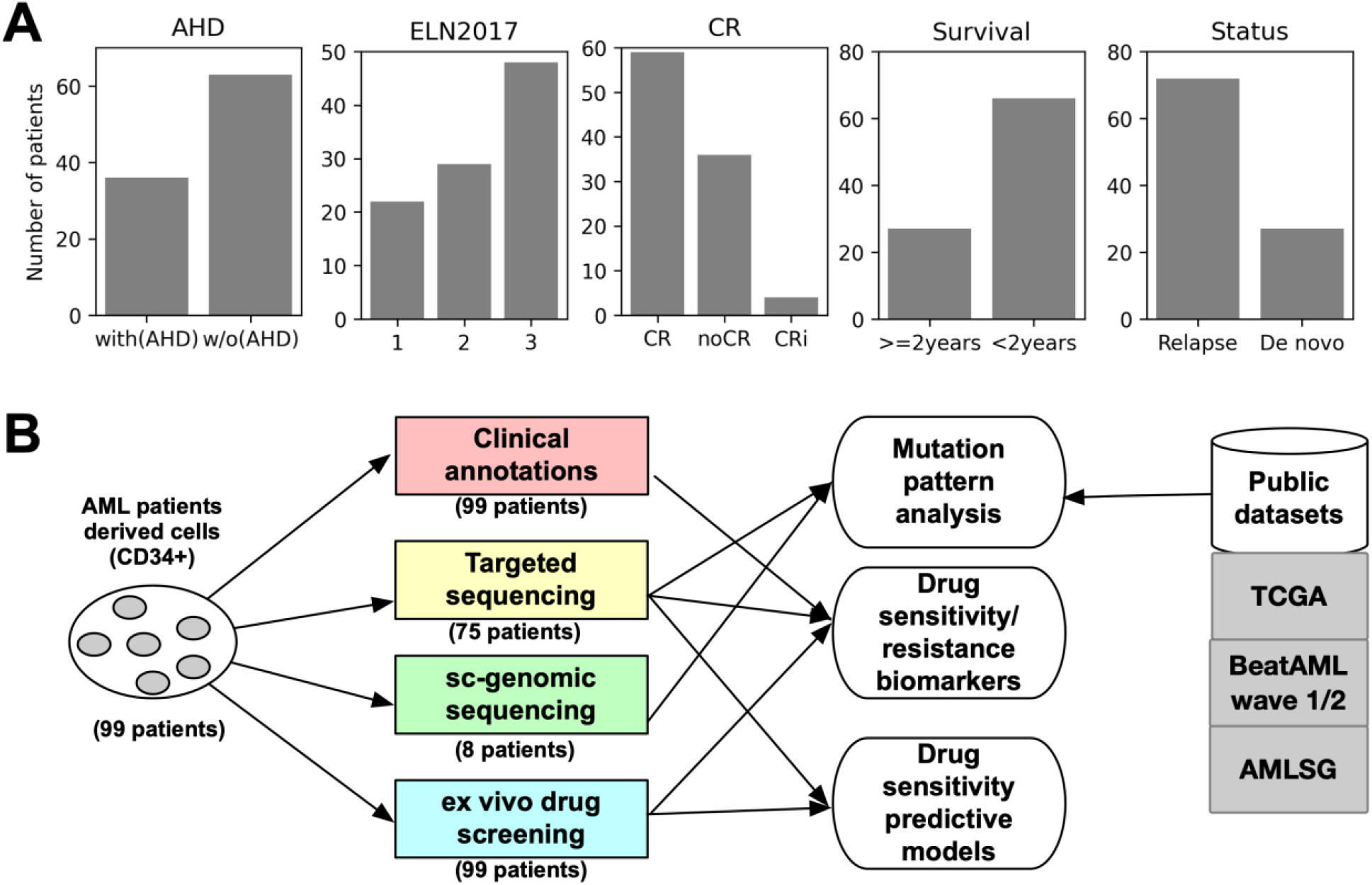
Overview of the study. A. Distribution of the cohort in this study, including distribution of patients with/without antecedent hematologic disorder (AHD), distribution of patients in different European LeukemiaNet (ELN2017) classification, distribution of patients with/without complete remission (CR) or partial remission, distribution of patients with survival time either greater or less than 2 years, and distribution of patient status, relapse or de novo. (B) Overview of the data collection, experimental design and study workflow.

### Genetic variants for AML

Genetic variants for common AML-related genes were sequenced using the MyAML^®^

Gene Panel Assay, examining 194 genes, including 36 translocations with an average sequencing depth greater than 1000x, allowing the detection of variants with low allele frequency. Potential causal variants were detected by filtering the common variants from large published populations (ExAC (Lek et al., 2016) and gnomAD (Karczewski et al., 2020)). By excluding variants with population-based allele frequency >0.01 observed in ExAC and gnomAD, we found AML-related genes such as DNMT3A (with mutation frequency: 28%), WT1 (25%), NRAS (25%), NOTCH2 (24%), TET2 (23%), RUNX1 (21%), FLT3 (21%), STAG2 (19%), TP53 (17%), NPM1 (17%), PTPN11 (13%), NF1 (12%) and CEBPA (12%), which are among the most frequently mutated genes [Figure 2, Supplementary Table 3]. The mutation frequency distribution is similar to previous studies including TCGA (Cancer Genome Atlas Research et al., 2013), Beat AML (Tyner et al., 2018) and AMLSG (Gerstung et al., 2017; Papaemmanuil et al., 2016). The distribution of variant allele frequencies for the high frequently mutated genes and sites are shown in Figure S1. The alterations of NRAS and KRAS are shown with amino acid substitutions at positions 12 and 61 [Supplementary Figure S1]. The alterations in WT1, NPM1, and FLT3 were more relevant to deletions or insertions. The distribution of VAF for specific sides in several other highly frequently mutated genes are also showed in the Supplementary Figure S1.

**Figure 2.**
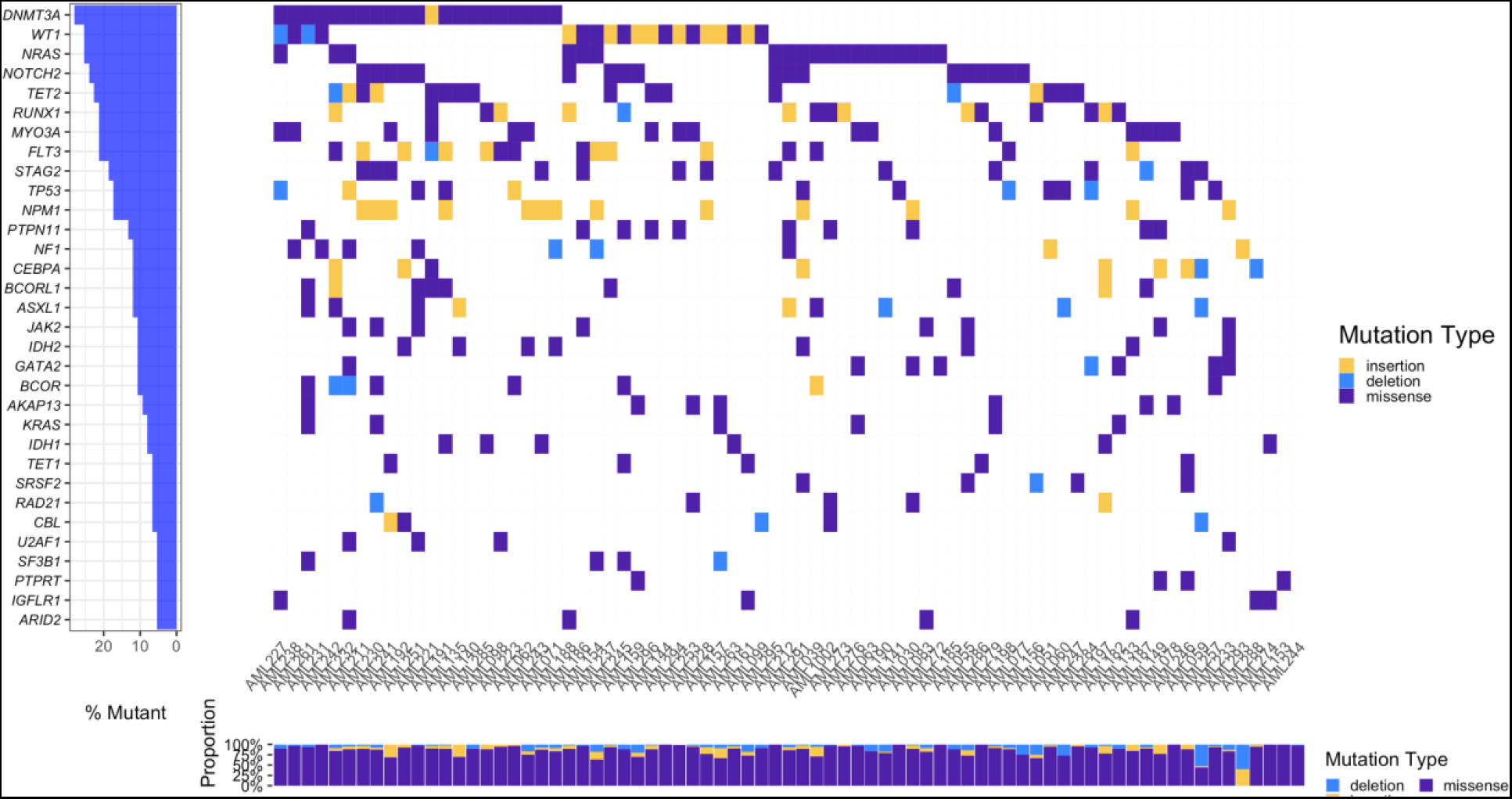
Genetic variants landscape for AML patients in the current study. Genes with mutation frequency greater than 5% of the tested samples, and are reported to be associated with AML are shown in the upper panel. Mutation types for each gene are colored using different colors.

### Drug screening expanded the selection of potential AML drugs

The *ex vivo* drug screening library includes 208 drugs or inhibitors that target a broad spectrum of pathways and molecular aberrations, including inhibitors that target *ABL*/*Src*, ALK, RTK, BCL- 2/BCL-xL, BET, CDKs, CHK, HDAC, DOT1L, hedgehog signaling, HSP90, IDH1, IDH2, IKK, IGF1R, MEK, MET, mTOR, PARP, PI3K, PKC, proteasome, RAF, and WNT signaling [Supplementary Table 4, the list of abbreviations can be found in Supplementary Table 5]. Among all the drugs tested, 98 drugs are FDA approved drugs and 37 drugs have been studied in clinical trials. The current FDA approved AML drugs include idarubicin, venetoclax, daunorubicin, azacitidine, cytarabine, decitabine, doxorubicin, enasidenib, ivosidenib, midostaurin, and mitoxantrone. Although there are immunotherapies that are FDA approved, including gemtuzumab ozogamicin and tagraxofusp (approved for blastic plasmacytoid dendritic cell neoplasm but in clinical trial for AML), these cannot be assayed in our current format due to 72h incubation and absence of immune cells given that there is enrichment of leukemic blasts. Log2-transformed IC50 (μM, inhibitory concentration at half maximal effect ([Supplementary Table 6] and area under the curve (AUC) values [Supplementary Table 7] are measured for each drug in different patients [see Methods].

The large drug screening library allows us to select potential drugs for AML patients according to the drug IC50. With the criteria of drugs having an IC50 less than 200 nM in at least 10% of the tested samples revealed 69 drugs that show potential utility in either most or some of the samples [Figure 3A, Supplementary Table 8]. Among them, we observed the FDA approved AML drugs idarubicin (with IC50 < 200nM in 96% samples), venetoclax (83%), mitoxantrone (83%), daunorubicin (81%), clofarabine (76%), ponatinib (45%), volasertib (27%). The percentage of samples with IC50 smaller than 200 nm for midostaurin, cytarabine and azacitidine is about 8%, 10% and 3%, respectively. Idarubicin, venetoclax, mitoxantrone, daunorubicin and clofarabine show higher sensitivity in more than 65% of the samples given the threshold selected for IC50. The drug concentration threshold for the selection of sensitive or resistant samples needs to be compared to the concentration achieved in patients to be clinically relevant. For example, high dose cytarabine has peak plasma concentration of 115 micromolar and 94% of patient samples are sensitive to cytarabine if the threshold is raised to 115 micromolar. Azacitidine and decitabine are hypomethylating agents, known to affect the methylation of the CpG islands that affect gene expression (Stomper et al., 2021), that don’t exhibit much direct cytotoxicity as single agents in the 72h assay and often take 1-6 months in patients to achieve remission as their action is related to changes in gene expression. There are alternatives to determining drug sensitivity to azacitidine (Notarstefano et al., 2021). The IDH inhibitors ivosidenib and enasidenib exhibit low drug sensitivity in the assay as they also do not have in vitro direct cytotoxicity as single agents due to their mechanism of action of suppressing the high levels of 2-hydroxyglutarate that occur in IDH mutated AML that block differentiation (Stubbins and Karsan, 2021).

**Figure 3.**
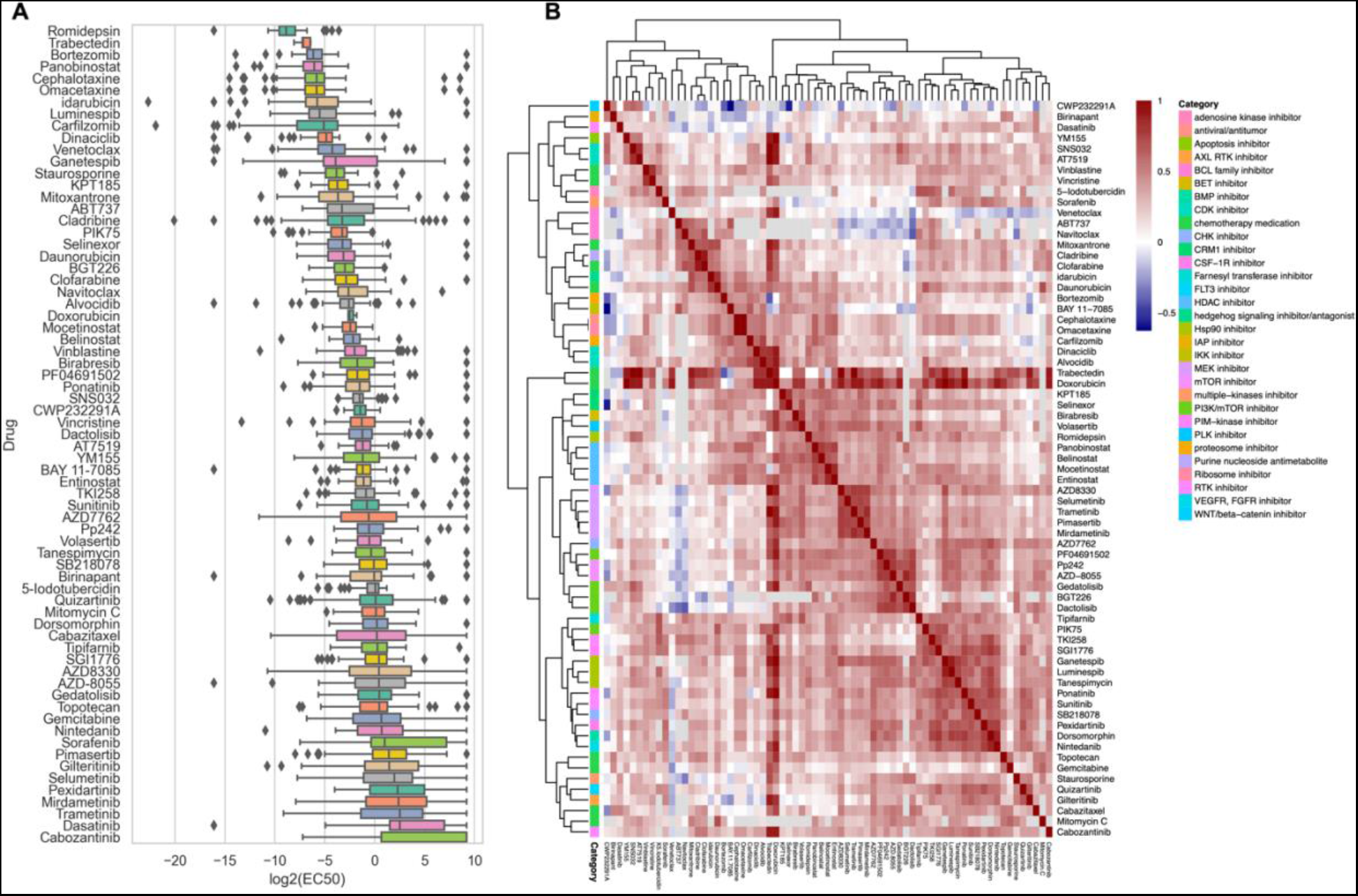
Drug sensitivity distribution and similarity. A. Boxplot of IC50 values for drugs with IC50 smaller than 200nM in at least 10% samples. X-axis: log2 transformed IC50 values. B. Hierarchical clustering of Spearman correlation coefficient for drugs shown in panel A.

Beyond the observation of different drug responses of FDA approved AML drugs, we also found other drugs or chemicals that show potential sensitivity in AML samples, including drugs that have been approved for the treatment of other diseases or undergoing clinical investigation [Supplementary Table 8]. 98% to 46% samples show sensitivity to the histone deacetylase (HDAC) inhibitors romidepsin (approved for multiple myeloma), panobinostat (under clinical trial for multiple myeloma), mocetinostat (under clinical trial for diffuse large B-Cell lymphoma), and belinostat (under clinical trial for multiple cancers). Most AML samples show sensitivity to the 20S proteasome inhibitors bortezomib (approved for multiple myeloma) and carfilzomib (approved for lymphoma), the HSP90 inhibitors luminespib and ganetespib, and the CDK inhibitors dinaciclib and alvocidib). More than half of the samples show sensitivity to the BCL-2 inhibitors venetoclax, navitoclax and ABT737. Over 75% samples show sensitivity to CRM1 inhibitors KPT185 and Selinexor. We also observed that 34% to 11% of the samples show sensitivity to the apoptosis inhibitor YM155 and the NF-kB inhibitor BAY 11-7085. Drugs such as MEK inhibitors, PI3k/mTOR inhibitors, VEGFR/FGFR inhibitors, CHK1 inhibitors, BMP inhibitors, JAK inhibitors and BET inhibitors show high activity in a smaller percentage of samples.

Hierarchical clustering of the Spearman correlation coefficient matrix for the drug sensitivity data reveals that drugs with similar actions or targets are often highly correlated, indicating a similar drug response. For example, the MEK inhibitors selumetinib, trametinib, pimasertib, AZD8330 and mirdametinib show similar response patterns [Figure 3B, Supplementary table 9]. The BCL family inhibitors venetoclax, navitoclax and ABT737 are also clustered together [Figure 3B].

### Drug sensitivity or resistance associated biomarkers

As many of the drugs show sensitivity in a subset of the AML samples, we investigated the genes or mutations that could serve as biomarkers for patient stratification for each of the drugs tested. We applied Rank-sum tests to test the significance of each gene that contains genetic variants in at least 5 samples by measuring the difference of median IC50 values in the samples with and without mutations in that gene. Among the top significant gene-drug sensitivity / resistance associations, we find the TP53 mutation is associated with resistance to AML drugs cytarabine and mitoxantrone *(P-*value *< 0.01)* [Figure 4A]. The mutations in CEBPA, ASXL2 and ELL are associated with sensitivity to venetoclax *(P-*value *< 0.01)*. The association between the CEBPA mutation and sensitivity to venetoclax can be also supported from a clinical trial observation that suggested the association of CEBPA biallelic mutation with a favorable response of venetoclax(Gangat and Tefferi, 2020). Our result also indicates the mutations in PRDM9, KMT2A, SRRM2, STAG2 are associated with drug sensitivity to daunorubicin *(P-*value *< 0.01)* and the mutation of SRRM2 is associated with sensitivity to azacitidine *(P-*value *< 0.01)*.

**Figure 4.**
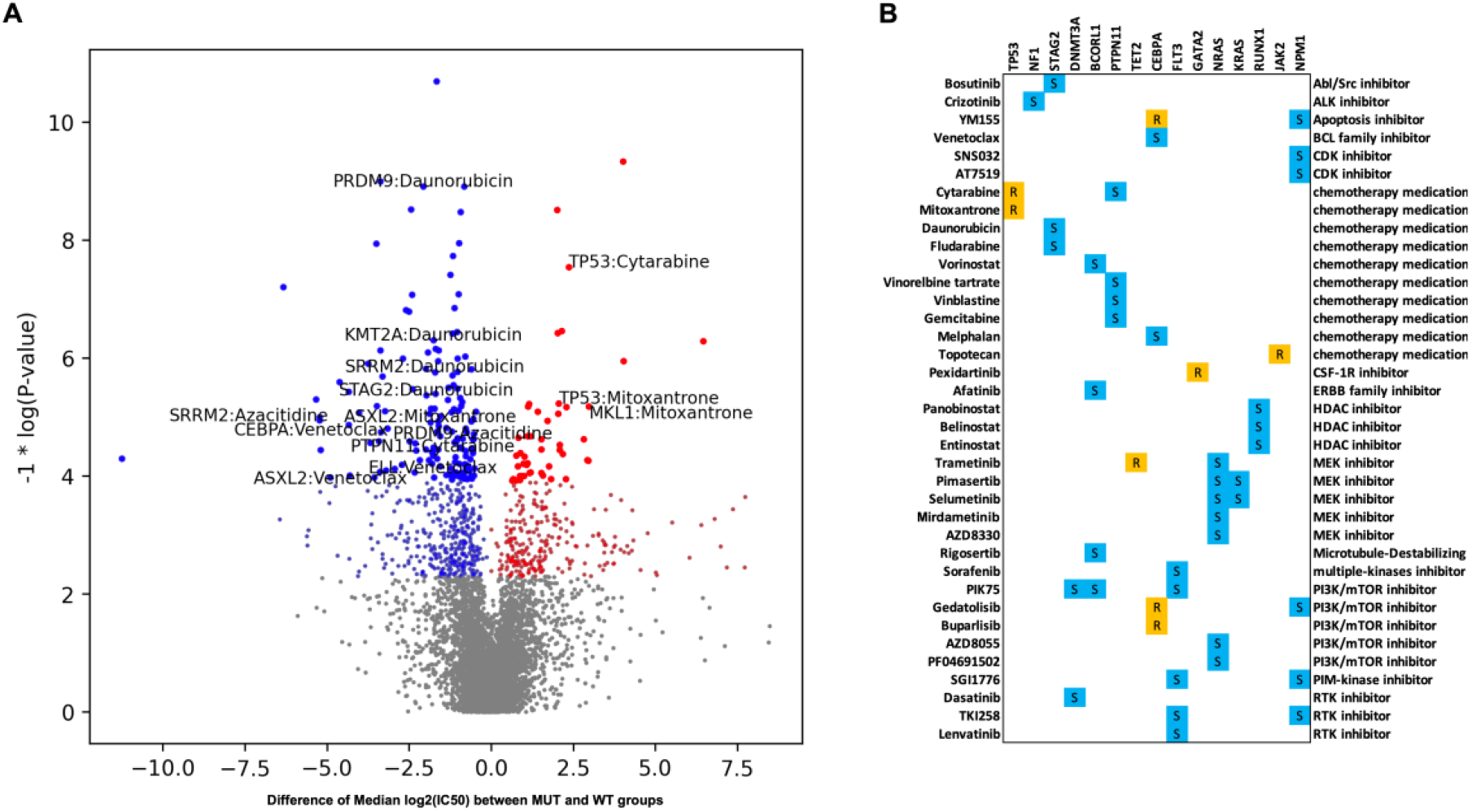
Gene signatures that are associated with drug response. A. Scatter plot for the gene - drug sensitivity / resistance associations. Red: association of resistance. Blue: association of sensitivity. The top significant gene - drug sensitivity / resistance association pair for the FDA approved AML drugs are labeled. The size of dots represents different significance levels: smaller dots represent the association is within the threshold of *P*-value < 0.05, and the bigger dots represent the association is within the threshold of *P*-value < 0.01. Y-axis: negative log transformed *P*-value in Rank sum test. X-axis: Difference of median log2 (IC50) for each drug between the mutated (MUT) group and the wild type (WT) group for each gene. B. Drug sensitivity associations with *P*-value < 0.01 for all the potential drugs and genes that are highly frequently mutated in AML. R: associated with resistance; S: associated with sensitivity.

The most frequently mutated genes in AML as observed in our datasets as well as other publications are associated with sensitivity or resistance to different types of drugs [Figure 4B]. The mutation of NRAS or KRAS is associated with a wide range of MEK inhibitors, including trametinib, pimasertib, selumetinib, mirdametinib, and AZD8330. Similar results are observed from the Beat AML dataset(Tyner et al., 2018). We also found that the mutation of NRAS is associated with sensitivity to PI3K/mTOR inhibitors AZD-8055 and PF04691502. TET2 mutation may be associated with resistance to Trametinib (*P-value < 0.01*). We also found that the mutation of TET2 is associated with resistance to a wide range of drugs, including mirdametinib, alvocidib, pimasertib, AZD8055, entinostat, paclitaxel, and birabresib. The genetic alterations of FLT3 are associated with sensitivity to receptor tyrosine kinase (RTK) inhibitors including sorafenib, PIK75, TKI258 and lenvatinib, and PIM-kinase inhibitor SGI1776 [Figure 4B]. Similarly, the mutation of NPM1 and DNMT3A is also associated with sensitivity to certain RTK inhibitors, such as TKI258 and dasatinib. The similar associations for different genes may be due to similar functionality such as NRAS and KRAS, or due to the co-occurrence of the gene mutations. We also found the RUNX1 mutation is associated with sensitivity to a BET inhibitor birabresib with *P-value* < 0.05 [Supplementary Table 10]. The TP53 mutation is associated with resistance to a wide range of drugs, including mitoxantrone, AMG900, cytarabine, clofarabine, cladribine, idarubicin, and melphalan; we also observed that it is associated with sensitivity to an NF-*κ*B inhibitor BAY11- 7082 [Supplementary table 10]. The power of single gene alterations for the association with drug sensitivity is low. When applying multiple correlation tests, none of these associations show significance with the threshold of FDR smaller than 0.05. This may be due to the limitation of sample size, but it also suggests that comprehensive gene mutation patterns in which mutations are aggregated on the basis of their co-occurrence or mutual exclusivity may work better for the prediction of drug response than single gene mutations.

### Co-occurring mutation patterns for AML

To attain a more systematic understanding of the mutation landscape of AML and how the genetic patterns are associated with drug sensitivity, we analyzed public datasets to learn the genetic mutation patterns in AML. The Beat AML(Tyner et al., 2018), TCGA-LAML(Cancer Genome Atlas Research et al., 2013), and AMLSG(Gerstung et al., 2017; Papaemmanuil et al., 2016) datasets share 15 genes in common that show mutation frequency in over 5% for AML patients, including FLT3, NPM1, DNMT3A, IDH2, RUNX1, IDH1, TET2, NRAS, TP53, CEBPA, WT1, PTPN11, KRAS, ASXL1, and STAG2. The highly frequently mutated genes show consistency across different projects as shown in [Supplementary Figure S2 A & B]. The mutations of the 15 commonly mutated genes dominate the AML samples from the three large cohort studies (Beat AML: 92%, TCGA: 79%, AMLSG: 87%) [Supplementary Figure S2 C]. To investigate the co-occurring and mutually exclusive patterns in AML patients, we used multiple statistical methods (Fisher exact test, permutation test, and DISCOVER(Canisius et al., 2016) on multiple datasets (TCGA-LAML, Beat AML, AMLSG or a combined dataset of TCGA-LAML and Beat AML), and integrated the statistical results using the rank aggregation methods(Kolde et al., 2012). We further transformed the co-mutation pairs into a co-mutation graph by assigning a weight for each co-mutation pair or mutually exclusive pair as a negative log10-transformed ranking aggregation score (adding 1 to the score to make sure the weight for the graph will be a positive number). The co-mutation graph is shown in Figure 5A and Supplementary Table 11.

**Figure 5.**
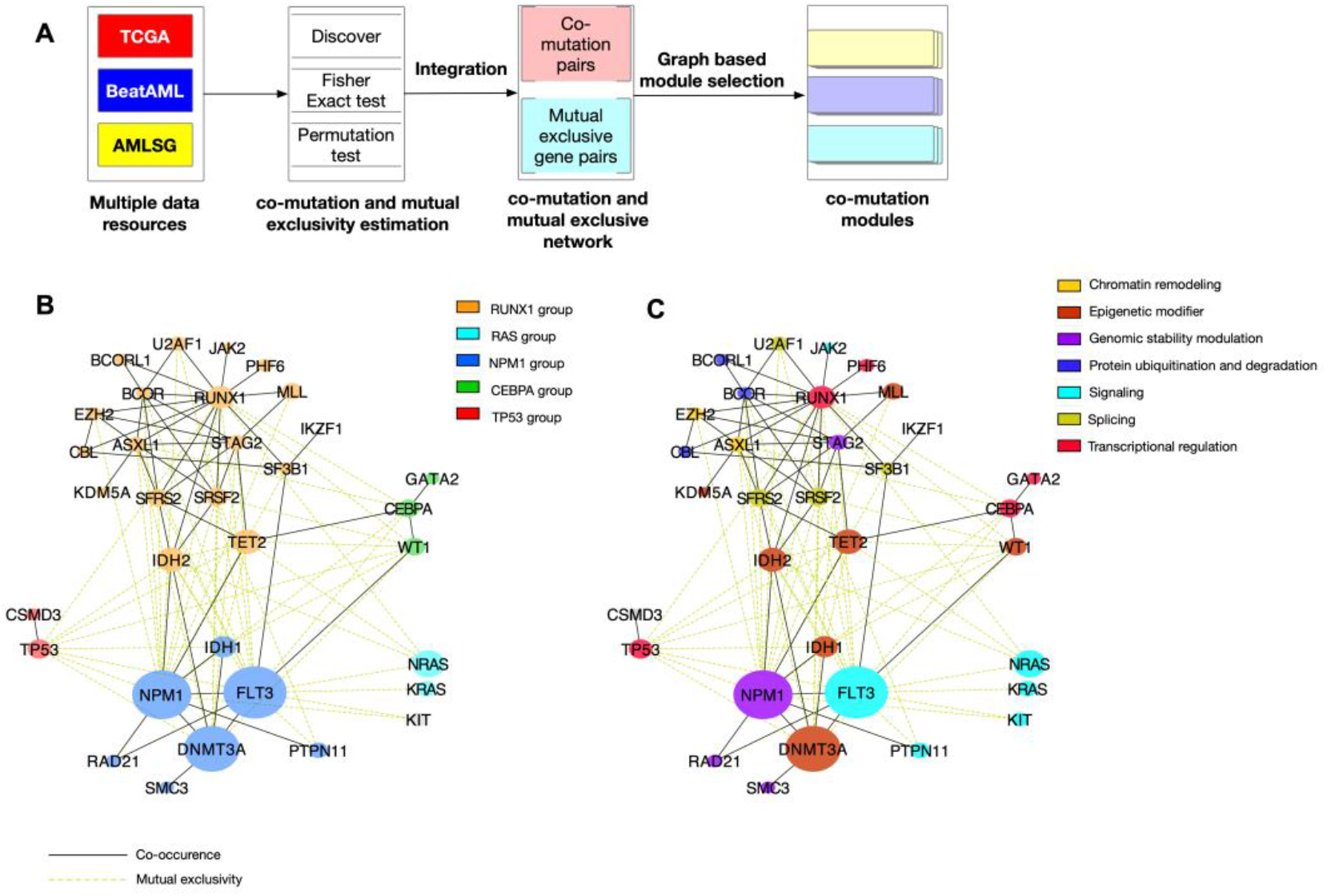
Co-occurring mutation clusters were revealed from the weighted co-mutation graph from multiple AML cohorts. A. An overview of workflows of detection of co-mutation modules, incorporating multiple data sources and methods to identify co-mutation and mutually exclusive gene pairs. Additionally, graph-based approaches were utilized to detect sub-co-mutation graphs. B. Within the co-mutation graph, five sub-graphs have been defined. C. The genes within different sub-co-mutation graphs are categorized into specific groups, allowing for a better understanding of their functional impacts.

To have a more systematic understanding of the mutation patterns, we used a graph-based approach (Pons and Latapy, 2006) to investigate the community of the mutated genes [Figure 5A]. The NRAS and KRAS mutations are not covered by the co-mutation graph, but they show high mutation frequency (16%) in AML(Padmakumar et al., 2021). We then added NRAS and KRAS to the graph as an independent group. As shown in Figure 5B, five co-occurring mutation groups were defined based on the analysis of a random walk based on subgraph detection (cluster_walktrap function in igraph package). We named the five groups as RUNX1 group, CEBPA group, NPM1 group, TP53 group as these key genes have the highest degrees in the co-mutation sub-graph, and the remaining RAS group includes only RAS (NRAS, KRAS) mutations. To confirm the co-occurring mutation clusters, we also applied another algorithm, Be-With (Dao et al., 2017), which considers both the co-mutation and the mutual exclusivity of mutations to obtain the clusters that are enriched in co-mutated genes in each cluster, but have mutually exclusive mutations in different clusters. This method gives similar results, but doesn’t include all of the genes. The results from the Be-With algorithm show NPM1, FLT3, DNMT3A and IDH1 are in one cluster, RUNX1, SRSF2, STAG2, ASXL1, IDH2, BCOR and SFRS2 are in another cluster. TET2 is clustered with CEBPA and WT1 using the Be-With algorithm as it has a mutual exclusivity relationship with the other genes in the RUNX1 cluster. The TET2 gene exhibits mutual exclusivity with IDH1, IDH2, NRAS, WT1, and TP53 genes. However, it shows co-occurrence with CEBPA, NPM1, SFRS2 and STAG2 genes.

By grouping the highly frequently mutated genes into different functional processes with which they are associated, we found that different co-mutation sub-graphs show different features [Figure 5C]. The RUNX1 co-mutation group is featured with the alteration of epigenetic modifiers and splicing factors. The RUNX1 group shows alterations in a larger number of genes, including the chromatin remodeling factors ASXL1, EZH2, transcription factor RUNX1, the splicing factors U2AF1, SFRS2 and SRSF2, the epigenetic modulators TET2, IDH2, KDM5A, MLL, the protein ubiquitination and degradation factors BCOR and BCORL1, and PHF6 [Figure 5C]. The TP53 group and CEBPA groups are dominated by dysregulation of different transcription factors. The CEBPA group includes mutations of CEBPA, GATA2, and WT1. Patients with gene mutations in this group are associated with abnormal transcriptional regulation and epigenetic dysregulation. The NPM1 co-mutation group features alterations in different factors, including the epigenetic modifier DNMT3A, the genomic stability modulator NPM1, kinase FLT3, and the DNA damage-related gene RAD21. The mutation of FLT3 alters the PI3K-RAS-MAPK signaling pathway. The KRAS and NRAS mutations, in the RAS group, also affect the RAS/MAPK signaling pathway, but downstream of FLT3.

We observed that the genes that play similar roles in a signaling pathway show mutual exclusivity, such as KIT and FLT3; both of them are receptor tyrosine kinases, which initiate the signaling in MAPK pathways. Our results show that the FLT3 and PTPN11 belong to the NPM1 co-mutation group. However, each of them shows co-occurrence with NPM1, yet mutual exclusivity is observed between FLT3 and PTPN11 (Figure 5C). IDH1 and IDH2 show mutual exclusivity, but show co-occurrences with the RUNX1 group and NPM1 group respectively. An independent cohort of 60 AML patients also shows the most frequently found mutations that co-occurred with IDH1/2 mutations were DNMT3A, SRSF2, ASXL1, and RUNX1(Wang et al., 2021).

The co-occurrence of gene mutations could exist at the patient level or at the single-cell level. At the single-cell level, the co-occurrence of gene mutations arises from the clonal hierarchy during the evolutionary history of the tumor’s development, while at the patient level, the co-occurrence of gene mutations results from the coexistence of two sub-clones(Morita et al., 2020). We hypothesize that the single-cell level co-occurrence may have a higher probability of co-occurrence, as the double mutations may contribute to cancer cell proliferation; on the other hand, the patient level coexistence of sub-clones may show a lower probability of co-occurrence as it would be more random. From the aggregated co-mutation graph (Supplementary Table 11), we found the co-mutation of FLT3-NPM1, DNMT3A-NPM1, DNMT3A-FLT3 are among the top of the co-mutated gene pairs, which is consistent with many other previous studies. Single-cell genotyping studies have proved that the three genes often co-occur in the same AML cancer cell(Morita et al., 2020; van Galen et al., 2019). Previous single cell genotyping studies also reported the coexistence of FLT3, NRAS, KRAS, PTPN11, and KIT in the same patient while they were often mutually exclusive at the cellular level. It is suggested that the sequence of mutation acquisition of distinct patterns of clonal evolution in AML follows linear and branching models of evolution(Morita et al., 2020).

### Clonal distribution at the single cell level

To understand the clonal heterogeneity of the tested patient samples, we performed single-cell genomic sequencing and simultaneous phenotyping using cell surface markers for selected patients using the MissionBio platform. We focused on eight representative samples (corresponding to the reagent kits that provide materials for eight samples) with multiple AML related genetic alterations for these single cell genetic analyses. We detected specific clones with distinct mutations using the MissionBio 45-gene myeloid panel. The analysis of the phenotype for these samples with the Biolegend TotalseqTM D Heme Oncology cocktail for 42 surface antigens demonstrated heterogeneity at the single cell level, but no differences in surface markers by subclone as detected in the eight samples. The variant allele frequency as measured in the bulk genome sequencing (NGS by MyAML®) is consistent with the variant allele frequency estimated using the single cell DNA sequencing (Figure 6A). The number of cells detected ranges from 4167 to 28723, with total reads ranging from 294 million to 893 million. Many subclones can be detected using the MissionBio gene panel. However, we detected some major clones that constitute a large proportion of cell populations (Figure 6B). By defining cells using selected variants (Supplementary Table 12), different subclones can be observed for each patient derived sample (Figure 6C). Although multiple subclones exist in the same patient, we observe dominant clones exhibiting co-occurring variants in the same cells. In the bulk targeted sequencing, we observed mutual exclusivity of FLT3 and NRAS; however, in some patients, such as AML186, we can still observe that both variants exist in the same patient. By single cell genome sequencing we observed cells with a FLT3 mutation or an NRAS mutation are commonly found in different cells, or different clones, which still show mutual exclusivity at the single cell level. As different clones may respond differently to drugs, it will be important to consider the co-occurrence of mutations within clones and the relative proportion of clones to provide better predictions for drug response.

**Figure 6.**
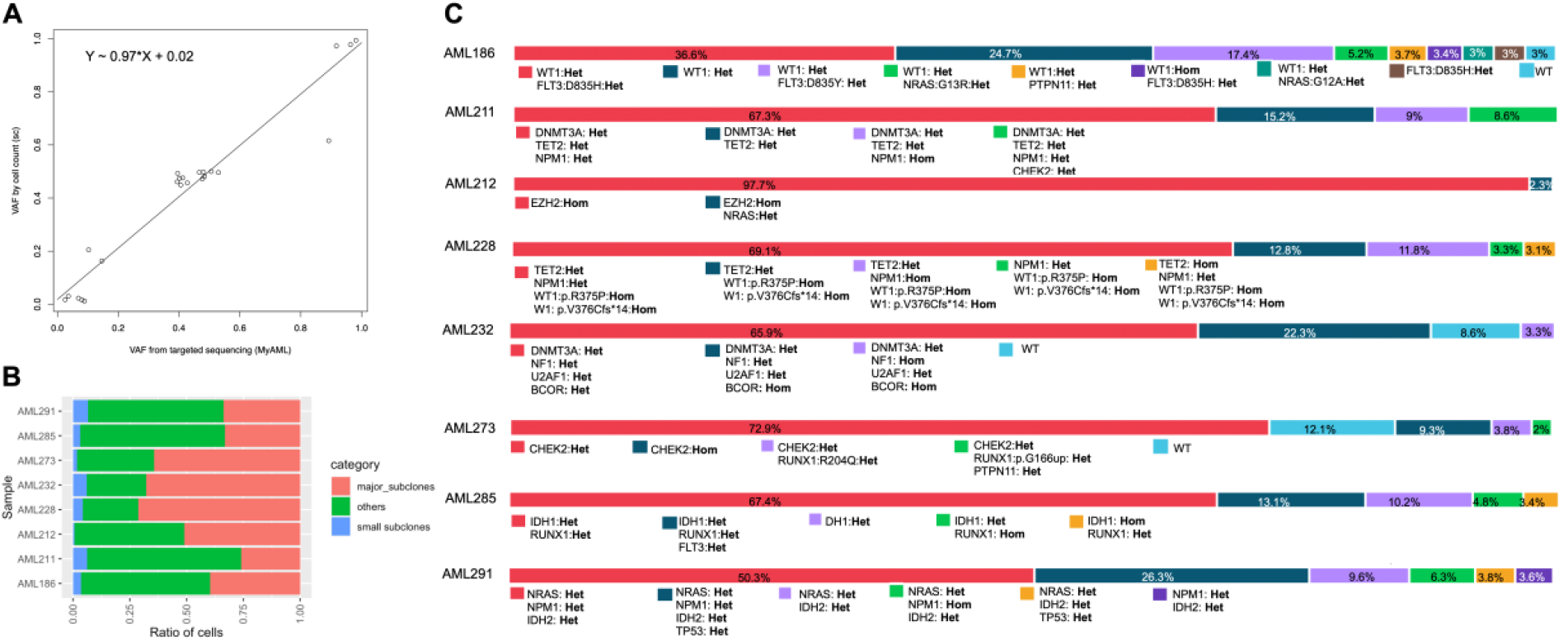
Sub-clones from the single cell genomic sequencing. A. Comparison between variant allele frequency (VAF) from the targeted sequencing (MyAML panel) and VAF estimated by cell count in the single cell experiment (Mission Bio). The lm function in the R package was used to model the relationship between the two variables. Data for the plot can be found in the Supplementary Table 12. B. Major, minor and other clones that can be explained by the selected variants. The variants selected for the subclone identification are listed in Supplementary Table 12. Here, major clones are defined by the subset of cells with the same genotype defined by the selected variants with a population of at least 1%. Minor clones are defined as the subset of clones with population smaller than 1%. Others are the cells missing the genotypes. C. Sub- clones defined by the selected variants for each patient derived sample.

### Prediction model using gene mutation patterns

We investigated whether the gene mutation patterns can predict drug sensitivity for different drugs. We selected drugs that have greater than 40 measurements for predictive model construction. Random Forest classifiers were used to train the prediction models for each drug. For the classification categories, we classified the samples into a drug sensitive group and a resistant group using either the 50% of the maximum plasma drug concentration or the median of the IC50, whichever was smaller. A stratified sampling approach was used to select a training set and an independent test set from both the resistant group and the sensitive group, with a ratio of 4:1. For the features, we considered the mutation of a single gene with mutation frequency greater than 3%, the co-occurrence pattern and the variant allele frequency as features to build prediction models. Hyperparameters were optimized using leave-one-out cross-validation in the training set using the GridSearchCV approach(Bergstra and Benjio, 2012). By randomly selecting 20 training sets and test sets, we evaluated the prediction accuracy on the test sets. The results suggest that gene mutation patterns are more predictive for some of the drugs or chemicals, such as the MEK inhibitors trametinib, selumetinib, and mirdametinib, chemotherapy drugs daunorubicin and mitoxantrone, CDK inhibitor alvocidib, FLT3 inhibitors quizartinib and gilteritinib etc [Figure 7A].

**Figure 7.**
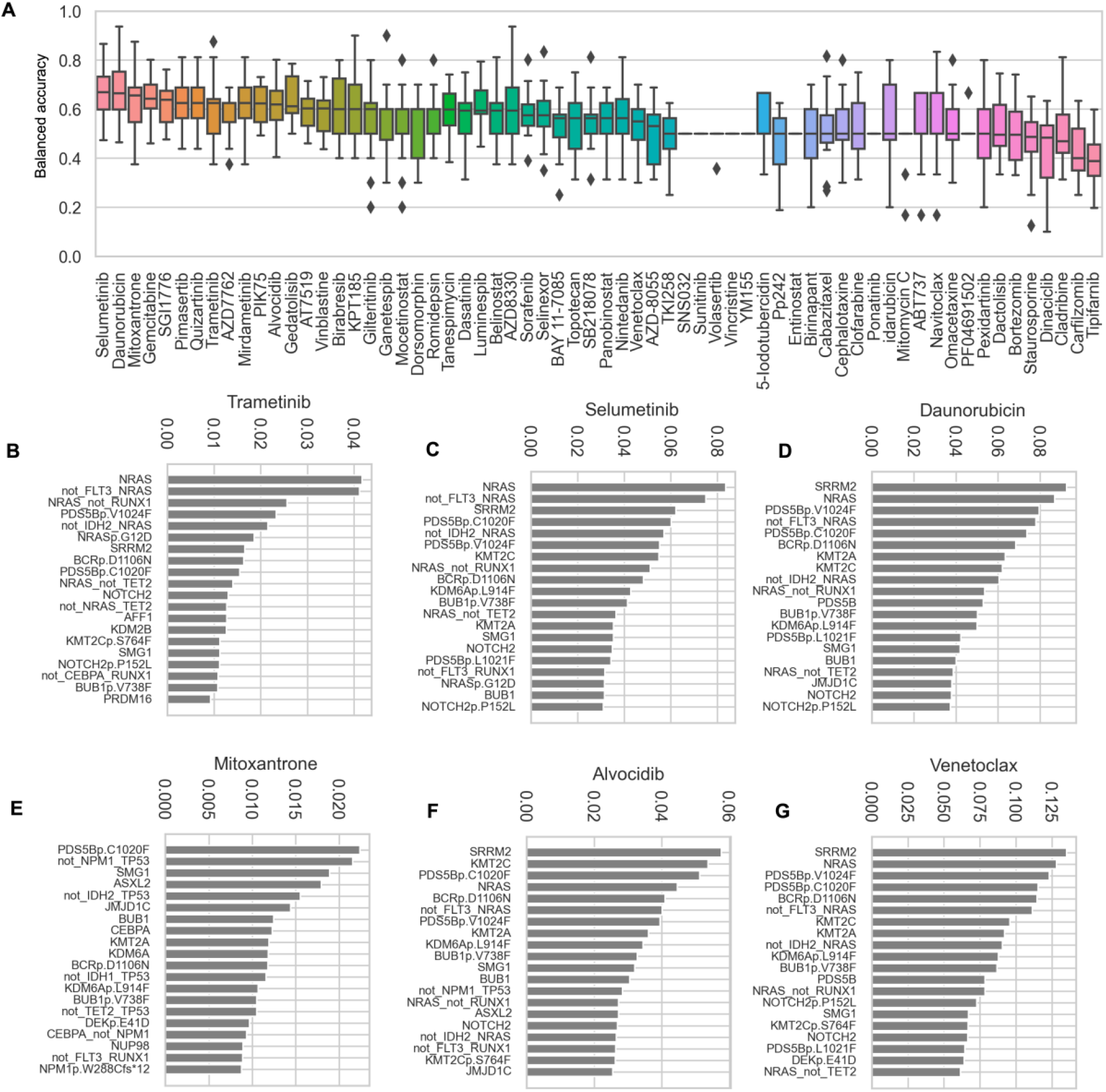
Prediction accuracy and most important features for the drugs from the Random Forest Models. A. Balanced accuracy in the testing datasets (20% from all samples) that were randomly selected. The models were first trained in the remaining 80% samples using leave one out cross-validation. B-G. Features with the most importance in the predictive models for selected drugs: Trametinib(B), Selumetinib (C), Daunorubicin(D), Mitoxantrone(E), Alvocidib(F), and Venetoclax (G). Top 20 features with average feature importance score from the Random Forest models with balanced accuracy score greater than 0.6 were shown. X-axis: Average feature importance.

We then selected models with balanced accuracy greater than 0.6 in the test sets to evaluate the most important features for the drug sensitivity prediction. The most important features for several selected drugs are shown in Figure 7B-G. The combinations of features such as FLT3-wildtype- and-NRAS-mutant or the allele frequency of NRASp.G12D are predictive of sensitivity to the MEK inhibitor trametinib [Figure 7B]. This is confirmed from the Spearman correlation between the genetic alteration events or their combinations and the drug response data [Supplementary Figure S3]. We computed Spearman correlation between the features and the log2 transformed IC50 values and performed BH-adjustments to adjust the *P*-values, and a threshold FDR < 0.1 were selected to highlight the features that are highly correlated with drug sensitivity (Supplementary Figure S3). The top important features for drugs that are in clinical application are shown in Figure 7C - G. The association between all the drugs in Figure 7A and genetic features can be found in Supplementary Table 13. The NPM1-wildtype-and-SRSF2-mutant is associated with resistance to daunorubicin (Supplementary Figure S3C). TP53 mutation and U2AF1 mutation are associated with resistance to mitoxantrone (Supplementary Figure S3D). The combinations of features, such as NPM1-wildtype-and-TP53-mutant, IDH2-wildtype-and-TP53-mutant, IDH1-wildtype-and- TP53-mutant, NPM1-wildtype-and-U2AF1-mutant are associated with resistance to mitoxantrone (Supplementary Figure S3D). KMT2A mutation is associated with sensitivity to alvocidib (Supplementary Figure S3E). We didn’t find variants that significantly correlated with the sensitivity of venetoclax using the threshold we chose (Supplementary Figure S3F). It may suggest other features still need to be taken into consideration when predicting the drug sensitivity of venetoclax, such as data at the expression level.

### *Ex vivo* drug screening predicts potential drugs for relapsed AML

Our *ex vivo* drug screening included both relapsed AML (72 samples) and de novo AML patients (27 samples). We then identified the drugs that show higher sensitivity in relapsed AML than in de novo AML patients. We found relapsed AML patients show higher sensitivity to several drugs, including arsenic trioxide (ATO), melphalan, rigosertib, afatinib, navitoclax and azacitidine [Figure 8A]. The samples with top ATO sensitivities exhibit either FLT3 mutations or TP53 mutations. Further clinical trials may be needed to validate this relationship. We also observed that navitoclax shows significantly higher sensitivity in relapsed AML patients [Figure 8A]. A clinical study has shown that navitoclax monotherapy has an acceptable safety and meaningful clinical activity in a minority of patients with relapsed/refractory lymphoid malignancies(de Vos et al., 2021). We further compared the drug response for patients with mutation in the same gene between the relapse and de novo patients. Among the patients with FLT3 mutation, we found drugs such as afatinib show higher sensitivity in the group of samples from the relapsed patients [Figure 8B]. On the other hand, dactolisib shows higher sensitivity in the de novo patient derived samples. Pazopanib has been approved for the treatment of patients with advanced renal cell carcinoma. Phase II clinical trial of pazopanib has been conducted in patients with AML(Kessler et al., 2019).

**Figure 8.**
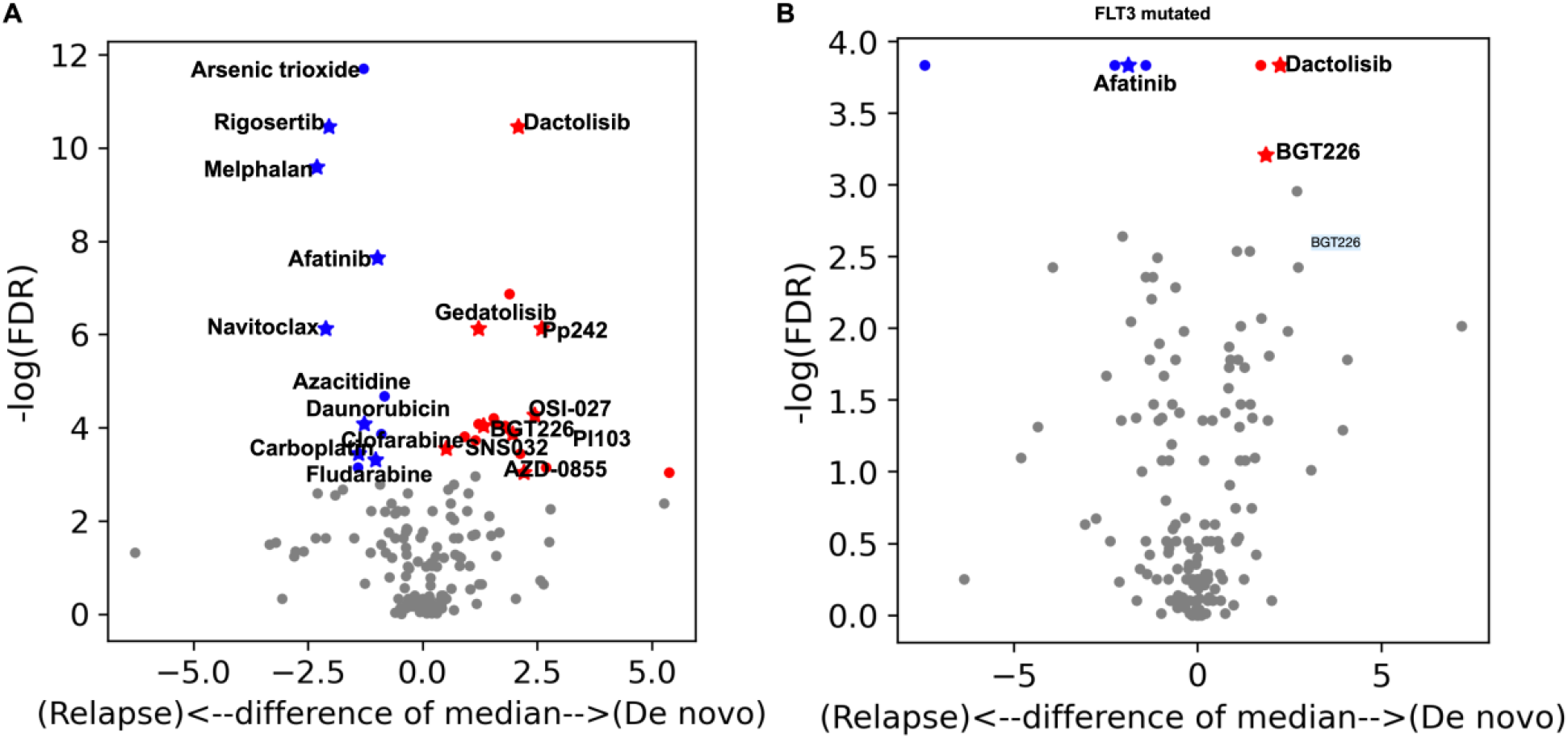
Differential response of drugs between the groups of relapsed and de novo AML patients with different mutation patterns. Rank sum test followed by BH adjustment was performed to estimate the significance of difference of IC50 values for each drug between the samples derived from relapsed AML patients and de novo AML patients. Differences of the median IC50 values for each drug between the relapsed group and the de novo group are used to estimate the difference. Drugs that show significant differences with FDR < 0.05 were highlighted in blue or red colors. Drugs colored blue are the ones that show significantly higher sensitivity in the relapse samples, and the drugs colored red are the ones that show significantly higher sensitivity in the de novo samples. Stars highlight the drugs with median IC50 smaller than 1µM in either the relapse samples or de novo samples. Drugs treated with sample size for both the de novo group and relapse group greater or equal to six were considered.

Our results suggest it shows higher sensitivity in relapsed AML patients, which may guide further clinical trial studies.

## Discussion

Our study provides a rich resource including the genetic variants and *ex vivo* drug screening data for more than 200 drugs for community use and provides clues for the selection of potential drugs or drug targets for further anti-AML drug development. The statistical associations between *ex vivo* drug screening and genetic information reveals biomarkers for the prediction of drug sensitivity or resistance for a variety of drugs. We have found that AML samples with RAS mutations show higher sensitivity to MEK inhibitors. We expect that the systematic analysis of the genetic patterns and definitions of co-mutation groups will facilitate more robust prediction of drug sensitivity or resistance.

The cohort in our study has a higher fraction of relapsed patients than other large datasets, such as TCGA-LAML and Beat AML. Our results suggested several drugs could have better response in the relapsed patients than in the de novo AML patients, providing clues to find more effective drugs for relapsed AML.

It is important to note that all the data for drug sensitivity and resistance reported here as well as in the other databases were based on *ex vivo* studies. Thus, the clinical utility of the predictions will require clinical trials. Furthermore, the *ex vivo* drug screening uses cell viability to quantify the efficacy of drugs, which may not reflect the effect of drugs that induce apoptosis or differentiation. New methodology with high throughput flow cytometry analysis will permit future studies to monitor these parameters. Lastly, for the inhouse studies, the target population was enriched with immunomagnetic bead separation, removing stromal cells and immune cells that are known to play a role in drug resistance. Some drugs, such as bortezomib, show high sensitivity in the *ex vivo* drug screening, although they may not be able to work well in patients. Further experiments or clinical trials are needed to validate the in vivo drug response effect.

## Material and Methods

### Study design

Samples from blood or bone marrow were collected from 99 AML patients with clinical annotation. Targeted gene sequencing and *ex vivo* drug screening were carried out to investigate the drug sensitivity- or resistance-associated biomarkers and to build the prediction models using the targeted sequencing assay. Integrative analysis from large datasets was carried out to characterize the gene co-occurrence patterns, which we further utilized as features for drug sensitivity prediction models.

### Sample collection and enrichment

Peripheral blood or bone marrow mononuclear cells were isolated from AML patients by density gradient centrifugation using lymphocyte separation medium. Blasts were enriched to >80% if needed by either positive selection for CD34, or depletion of non-myeloid populations using magnetic beads. For CD34 positive AML samples, cells were treated with FcR-blocking reagent and human CD34 Microbeads (Miltenyi Biotec, Gaithersburg, MD) for 30 min. Then, the magnetically labeled cells were enriched on positive selection columns inside the magnetic field of the MiniMACS. Non-target cells passed through the columns and were recovered in the elution buffer. After removing the columns from the MiniMACS separator, the retained CD34+ cells were eluted with a buffer using the plunger and recovered CD34+ cells were washed twice with culture medium. For the few AML samples that were CD34 negative, blasts were enriched by removing non myeloid populations as determined by clinical flow cytometry, most frequently by depletion of CD3+ lymphocytes and CD235a+ erythroid progenitors.

#### Drug response measurement and processing

The cell populations were analyzed for survival after a 72-hour exposure to 8-12 customized drug concentrations (within the range of 5pM to 100µM) of each drug spanning 4-5 logs. Post exposure viability was determined using CellTiter Glo luminescent reagent (Promega, Madison, WI) per manufacturer protocol, then the plates were analyzed with the Envision Multi-label plate reader (Perkin Elmer, Waltham, MA). XLFit (IDBS, Guildford, Surrey, United Kingdom) a Microsoft Excel Addin was used to analyze the data and generate dose response curves based on standard 4 - parameter logistic fit (i.e., fit = (A+(B/(1+((x/C)^D)))) where A and B equal minimum and maximum asymptotes, C equals IC50 and D equals slope). The Area under the curve values (AUCs) were calculated using the XLFit software utilizing minimum and maximum concentrations of drugs/compounds within the panel as the limits of the AUC calculation. For each plate, data were normalized to DMSO 100% viability and blank controls.

#### Targeted sequence analysis

MyAML™ uses next generation sequencing (NGS) to analyze the 3’ and 5’ UTR and exonic regions of 196 genes and potential genomic breakpoints within known somatic gene fusion breakpoints known to be associated with AML. Fragmented genomic DNA (∼3.4Mb) is captured with a customized probe design, and sequenced with 300bp paired end reads on an Illumina MiSeq instrument to an average depth of coverage >1000x. Using a custom bioinformatics pipeline, MyInformatics™, single nucleotide variants (SNVs), insertion/deletions (indels), inversions and translocations are identified, annotated, characterized, and allelic frequencies calculated. Commonly associated variants in dbSNP and 1000 genomes were eliminated. 3.4 Mb of DNA spanning the exons and a subset of introns from 194 genes were targeted by MyAML baits. 0.5-1.0 μg of patient DNA was sheared before hybridizing to MyAML baits. Captured target DNA was then sequenced using an Illumina platform.

A customized bioinformatics pipeline identified and characterized SNVs, indels, SVs, and CNVs. Average sequencing depth was greater than 1000x. More than 95% of targeted bases have at least 100x depth. The most commonly mutated genes in AML are all targeted with an average depth of coverage of 975x (range = 417x to 1370x). The overall analytical sensitivity is >96% and specificity >99.99% for SNVs. The overall analytical sensitivity is >95% and specificity >99.98% for indels. There is 100% sensitivity and 100% specificity for known pathogenic mutations, including missense and nonsense mutations in FLT3, DNMT3A, IDH1, IDH2, KIT, NRAS, KRAS, and TP53. The method accurately detects SNV and indels with allelic frequencies as low as 2.5% with >95% reproducibility.

#### Single Cell Mutation Analysis with Phenotype (DNA + Protein)

We utilized the MissionBio Tapestri® Platform and manufacturer provided protocols (Tapestri Single-Cell DNA + Protein Sequencing User Guide). We utilized the MissionBio 45-gene myeloid panel for targeted sequencing, with 312 amplicons, covering genes in myeloid disorders including AML, MDS, MPN, and CMML. Cryopreserved mononuclear cell samples from primary patient blood or bone marrow were thawed, incubated in IMDM with 15% fetal calf serum and 15% horse serum to enhance viability, then subjected to mononuclear cell isolation using lymphocyte separation media. Samples with less than 95% viable cells also underwent enrichment using the dead cell removal kit (Miltenyi). The viable cell fraction was verified by flow cytometry using viability dyes. The protocols provided by MissionBio for the analysis were followed to prepare DNA and protein libraries for sequencing. A total of 1 million cells were used for the initial steps of the procedure. The cells were incubated with Trustain FcX, then the TotalSeq™-D Human Heme Oncology Cocktail, V1.0 (Biolegend) panel of antibodies. Cells were washed several times and re-quantified to adjust the concentration to 3,000-4000 cells/mcl.A total volume of 35 mcl was loaded onto the Tapestri microfluidics DNA cartridge. Single cells were encapsulated, lysed, and barcoded. DNA and protein products were purified with AMPure XP beads and libraries prepared following MissionBio Tapestri user manual. DNA analysis and sequencing was performed by the University of California Irvine High Throughput DNA Sequencing Core (NCI Shared Resource) on an Agilent Bioanalyzer and Illumina NovaSeq 6000. 294 million to 893 million reads were obtained for each sample. The sequences were analyzed using Tapestri Insights software (MissionBio) to provide the characterization of the clonal mutations and distribution. The phenotype was analyzed using Mosaic software (MissionBio).

Initial steps for filtering low quality cells or genotypes were carried out in Tapestri Insights software with default parameters, which included removing genotype in cell with quality smaller than 30, genotype in cell with read depth smaller than 10, genotype in cell with alternative allele frequency smaller than 20, cells with smaller than 50% of genotypes present, and variants mutated in 1% of cells. We then selected known clinical variants that are annotated as “Pathogenic” or “Likely Pathogenic ” in the ClinVar database for subclone annotation or variants that have been reported to be in AML. The selected variants are shown in Supplementary Table 12. Subclones were identified using Tapestri Insights 2.2 using the selected variants. Major clones are defined by the subclones that show genotype in at least 1% of the cells. Allele drop outs were identified where there was a matching pair of a homozygous clone and wildtype clone.

#### Variants analysis

We filtered out all synonymous mutations, SNVs with minor allele frequency smaller than 2.5%, SNVs with population base minor allele frequency greater than 0.1% from the Genome Aggregation Database (gnomAD v2.1.1, reference genome GRCh37/hg19, both exon and whole genome-based datasets)(Karczewski et al., 2020) and ExAC database(Whiffin et al., 2017) (GRCh37/hg19 reference genome). GenVisR package was used to visualize the variants.

We further extracted mutations in genes that are highly frequently mutated in the TCGA-LAML(Cancer Genome Atlas Research et al., 2013), Beat AML(Tyner et al., 2018) and AMLSG (Gerstung et al., 2017; Papaemmanuil et al., 2016) projects, that include TP53, CSMD3, NPM1, FLT3, IDH1, DNMT3A, RAD21, SMC3, PTPN11, NRAS, KRAS, KIT, WT1, CEBPA, GATA2, TET2, IDH2, SFRS2, SRSF2, SF3B1, IKZF1, STAG2, ASXL1, KMD5A, CBL, EZH2, BCOR, BCORL1, U2AF1, JAK2, PHF6, MLL, RUNX1, NF1.

#### Co-mutation and mutual exclusivity analysis

Co-mutation and mutual exclusivity of gene mutations were estimated using three different methods, including Discover(Canisius et al., 2016), FisherExact test and Permutation test for four datasets separately, namely 1) TCGA-LAML(Cancer Genome Atlas Research et al., 2013), 2) Beat AML(Tyner et al., 2018), 3) targeted gene sequencing of 111 myeloid cancer genes from the German-Austrian AML study Group(Gerstung et al., 2017; Papaemmanuil et al., 2016), and a combined cohort from both TCGA-LAML and Beat AML. Genes that have been mutated in at least 5 samples were taken into account. All hypothesis tests were followed by Benjamini-Hochberg (BH) multiple testing correction. Co-mutation pairs were selected using the following criteria: 1) Gene pairs with false discovery rate (FDR) less than 0.05; and 2) the number of samples showing co-mutation greater than 3. Mutually exclusive gene pairs were selected based on FDR values (less than 0.05).

#### Aggregation of *P*-values

Ranked co-mutation or mutually exclusive gene pairs (FDR < 0.05) resulted from different resources (TCGA-LAML, Beat AML, AMLSG, combined cohort of TCGA & Beat AML) and methods were used to compute the aggregated score using the aggregateRanks function in the R package of RobustRankAggreg (Kolde et al., 2012). This algorithm allows to merge lists of different lengths. In the final list, it detects the items that are ranked consistently better than expected under the null hypothesis of uncorrelated inputs and assigns a significance score for each item.

#### Define Co-mutation patterns

Igraph was used to compute communities of co-mutated genes in the co-mutation graph using the random walk method (cluster_walktrap function in the igraph R package (Pons and Latapy, 2006). The co-mutation network is treated as a weighted undirected graph with weights for each node as 1-log10(aggregated score) resulting from the rank aggregation method.

BeWITH is used to identify modules with different combinations of mutation and interaction patterns(Dao et al., 2017). BeWITH algorithm has three different approaches (BeME-WithFun, BeME-WithCo and BeCo-WithMEFun) to find modules based on mutation co-occurrence, mutual exclusivity and functional interactions and it uses Integer Linear Programming (ILP) to find optimally scoring sets of modules. The approach we used in our project is BeME-WithCo. It Identifies modules which have co-occurring mutations within the modules while having mutual exclusivity between the modules. In its optimization function, it maximizes intercluster mutual exclusivity - co-mutation scores and in-cluster co-mutation - mutual exclusivity scores. The weight for the co-mutation and mutual exclusivity is defined weight (w)=max(1-log0p, 7) and the sensitivity is 10^-6 (any *P*-value smaller than 10^-6 considered as 10^-6) where p is the aggregated score resulting from the rank aggregation method.

### Prediction models using the gene mutation patterns

Random Forest classifier in sklearn python library is used to predict drug sensitivity using the mutation features, co-occurrence/mutual exclusivity features and variant allele frequency. We classified the samples into a drug sensitive group and a resistant group using either the 50% of the maximum plasma drug concentration or the median of the IC50 whichever is smaller. The maximum plasma drug concentration was collected from the literature. The dependent variables (features) include three categories: 1) the mutation status of genes which show mutation frequency of at least 3% (value range: 0 or 1), 2) the mutually exclusive pattern for genes in category 1 (0 or 1), and 3) the variant allele frequency for variants of genes in category 1. For each drug, a stratified sampling approach is used to select a training set and an independent testing set from both the resistant group and sensitive group, with a ratio of 4:1. Models are selected using leave-one-out cross-validation in the training set using the GridSearchCV function in the Sklearn python library, and are tested using the independent testing set using accuracy_score and balanced_accuracy_score function in sklearn.metrics package. The training process was repeated 20 times with randomly selected training sets after which the resulting models were used to predict the balanced accuracy scores for the remaining test sets. The median values of the balanced accuracy scores were used to rank the drugs [Figure 7A]. Models with balanced accuracy greater than 0.6 were selected to analyze the most important features that contribute to the prediction of drug sensitivity/resistance. For each drug, from all the models with a balanced accuracy score greater than 0.6, we averaged the feature importance and selected the top important features [Figure 7B-G].

We performed Spearman correlation analysis between all the feature and drug IC50 values. Multiple testing correction was performed using the BH method. Features with negative correlation suggest the observation of the event or high allele frequency of the variants is associated with sensitivity to the drug while features with a positive correlation suggest it is associated with resistance to the drug.

### Statistics

#### Correlation analysis of drug sensitivity data

Drugs with IC50 values smaller than 200nM in at least 10% samples were selected to perform correlation analysis. Spearman correlation was performed to measure the correlation coefficient between the IC50 values for each drug. Hierarchical clustering was used to cluster the correlation coefficient with ‘complete’ linkage and the clusters are visualized using pheatmap R package.

#### Gene mutation associated drug sensitivity analysis

We used Wilcoxon rank sum test to compare the drug IC50 value distribution between the samples with or without a gene mutation. Multiple testing correction was performed using the BH adjustment followed by Wilcoxon rank sum test. Genes with a *P*-value smaller than 0.05 in the rank sum test were designated as signature genes for drug sensitivity or resistance.

#### Clinical feature associated drug sensitivity analysis

We used the clinical feature of de novo or relapsed AML. We compared the drug IC50 values distribution between the two groups using Wilcoxon rank sum test followed by BH adjustment. Difference of the median log2 transformed IC50 values between the relapsed group and the de novo group were used to measure the difference.

### Data availability

All data are accessible from the supplementary tables.

## Acknowledgements

This project is supported by National Cancer Institute (NCI), National Institutes of Health (NIH) grants P01CA077852 (R.M.), and R01CA270210 (I.S.). This work was made possible, in part, through access to the Genomics Research and Technology Hub Shared Resource of the Cancer Center Support Grant (P30CA-062203) at the University of California, Irvine and NIH shared instrumentation grants 1S10RR025496-01, 1S10OD010794-01, and 1S10OD021718-01.

## Author contributions

Guangrong Qin led the computational analysis, performed statistical analysis and machine learning analysis, interpreted the results, drafted and revised the manuscript. Jin Dai and Sylvia Chien prepared the AML samples. Timothy Martins provides preliminary analysis for the drug sensitivity data. Brenda Loera analyzed single cell genomic sequence data. Quy Nguyen and Melanie Oakes analyzed the single cell mutation analysis library preps, provided advice and assistance, and performed and provided the sequencing data and description of the method. Bahar Tercan, Boris Aguilar performed statistical analysis for the gene co-mutation analysis. Lauren Hagen contributes to public data collection. Jeannine McCune provided the list of maximum plasma drug concentration. Richard Gelinas provided advice regarding the manuscript and revised the manuscript. Raymond J. Monnat, Jr inspired the research as PI of the Program Project grant that supported this effort, advised regarding development of the project, provided critical review of the progress, and revised the manuscript. Ilya Shmulevich supervised the project and revised the manuscript. Pamela S. Becker initiated the project, was PI of the IRB approved sample protocol, supervised the preparation of the patient samples for mutation analysis and high throughput drug screening, developed the methodology for the high throughput drug screening, conducted the single cell mutation analysis, interpreted the drug screening and single cell mutation data, reviewed the results of the NGS mutation analysis, drafted and revised the manuscript.

The authors declare no competing interests.

