## Supplemental figures for "Mutation Patterns Predict Drug Sensitivity in Acute Myeloid Leukemia"

**Figure S1. Variant allele frequency distribution of genetic variant events.** Variants are filtered using a population-based allele frequency threshold of 0.01 from ExAC and gnomAD datasets.

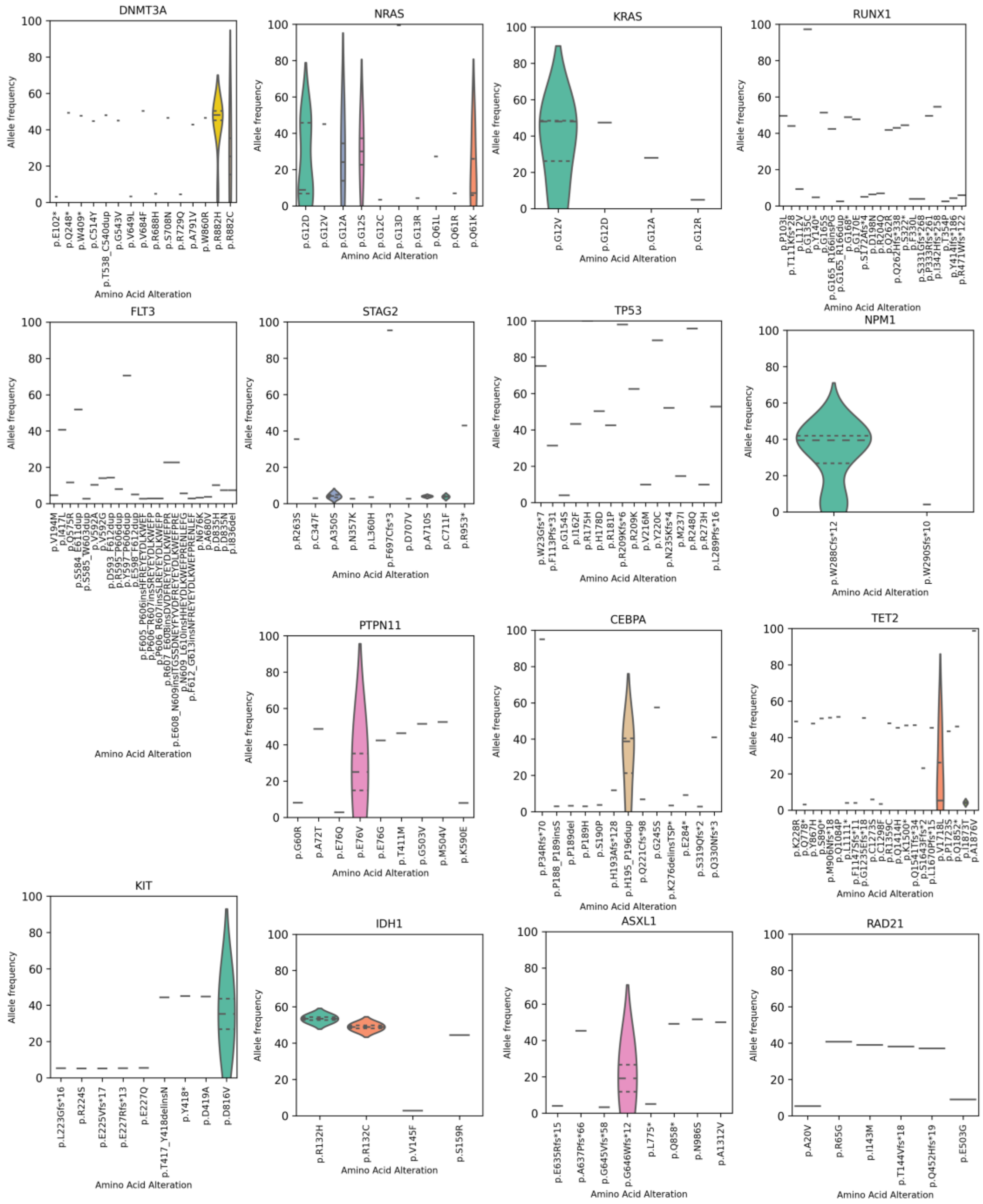

**Supplementary Figure S2**. Mutational landscape of AML from different projects. A. Highly frequently mutated genes in AML with frequency higher than 0.03 in any of the three projects, TCGA project (upper panel), AMLSG (middle panel), and the Beat AML project (lower panel). B. Overlap of the highly frequently mutated genes among three projects, only genes which show mutation frequencies higher than 0.05 were used. C. Coverage of samples with mutations among the top 15 genes.

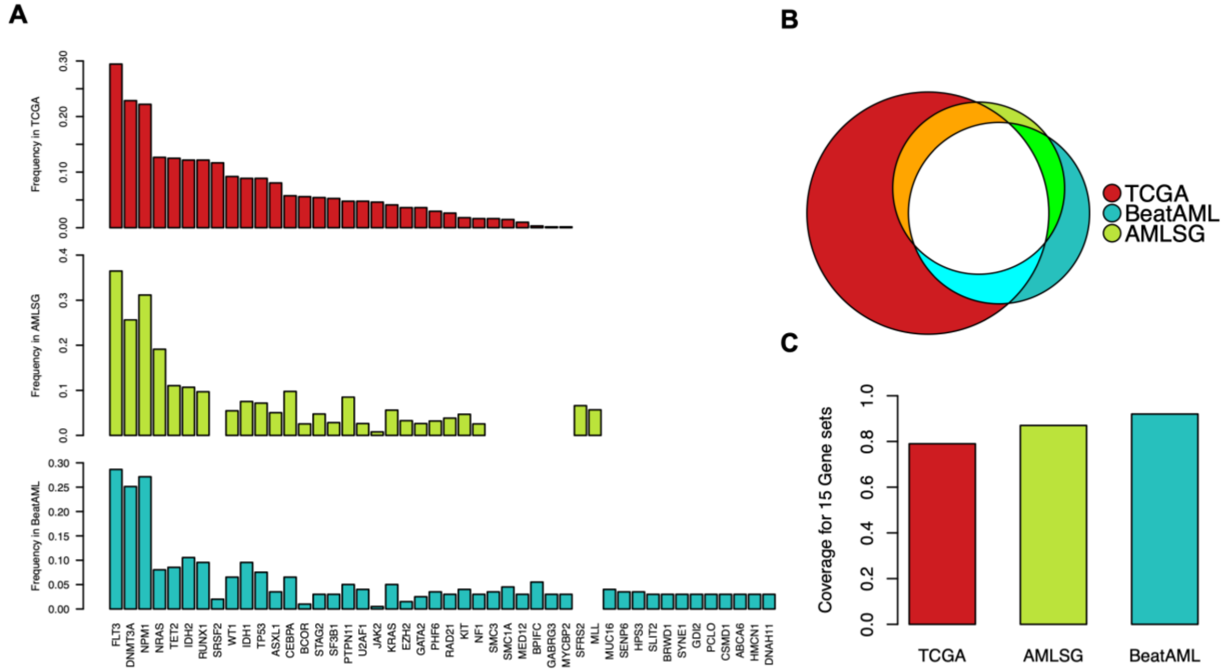

**Supplementary Figure S3.** Spearman correlations between the mutation features with drug IC50 values. X-axis represents the spearman correlation coefficient. Y-axis shows the negative log transformed FDRs using BH adjustment for the spearman correlation *P*-values. All labeled features are with FDR < 0.1. Blue color suggests the mutation of this gene or high variant frequency is associated with sensitivity of the drug. Red color suggests these features are associated with resistance of the drugs.

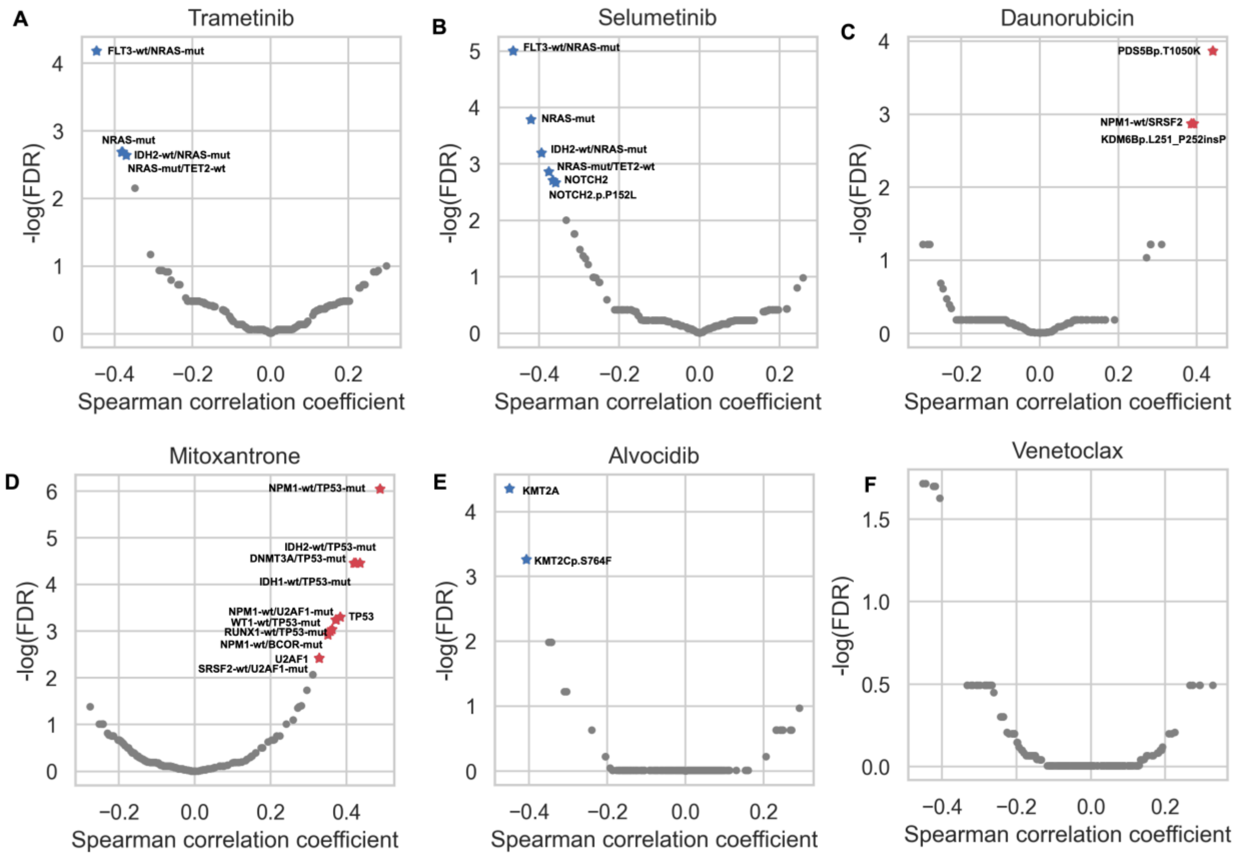
